## Supplemental Figures and Tables for "Adolescent length growth spurts in bonobos and other primates: Mind the scale"

Andreas Berghänel *et al.*

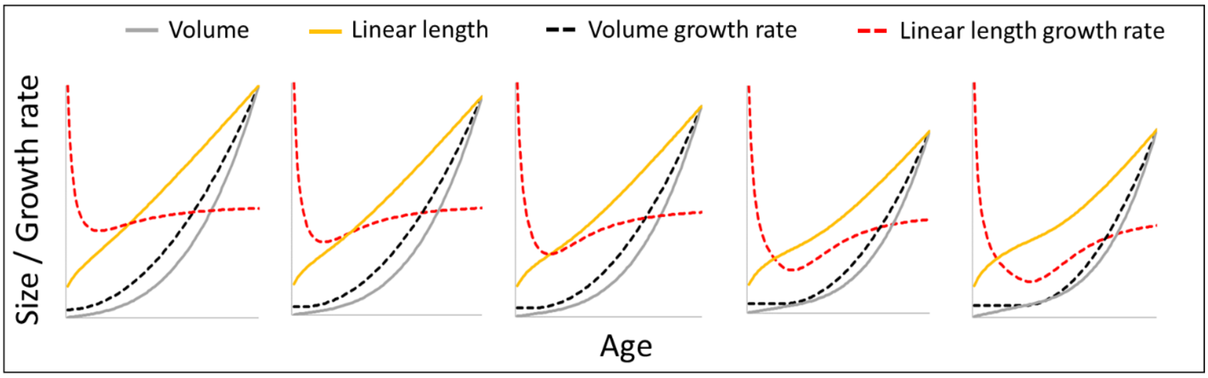


**Figure 1–figure supplement 1.** The figure shows how a change from a constant to a quadratically accelerating volume growth rate at a certain body size (identical levels of acceleration) results in different levels of acceleration in linear length growth rate depending on the age (in fact, body size) at which this change occurs, from left (change right after birth at small body size, only slight acceleration in linear length growth rate) to right (change at late age and larger body size, strong acceleration in length growth rate).

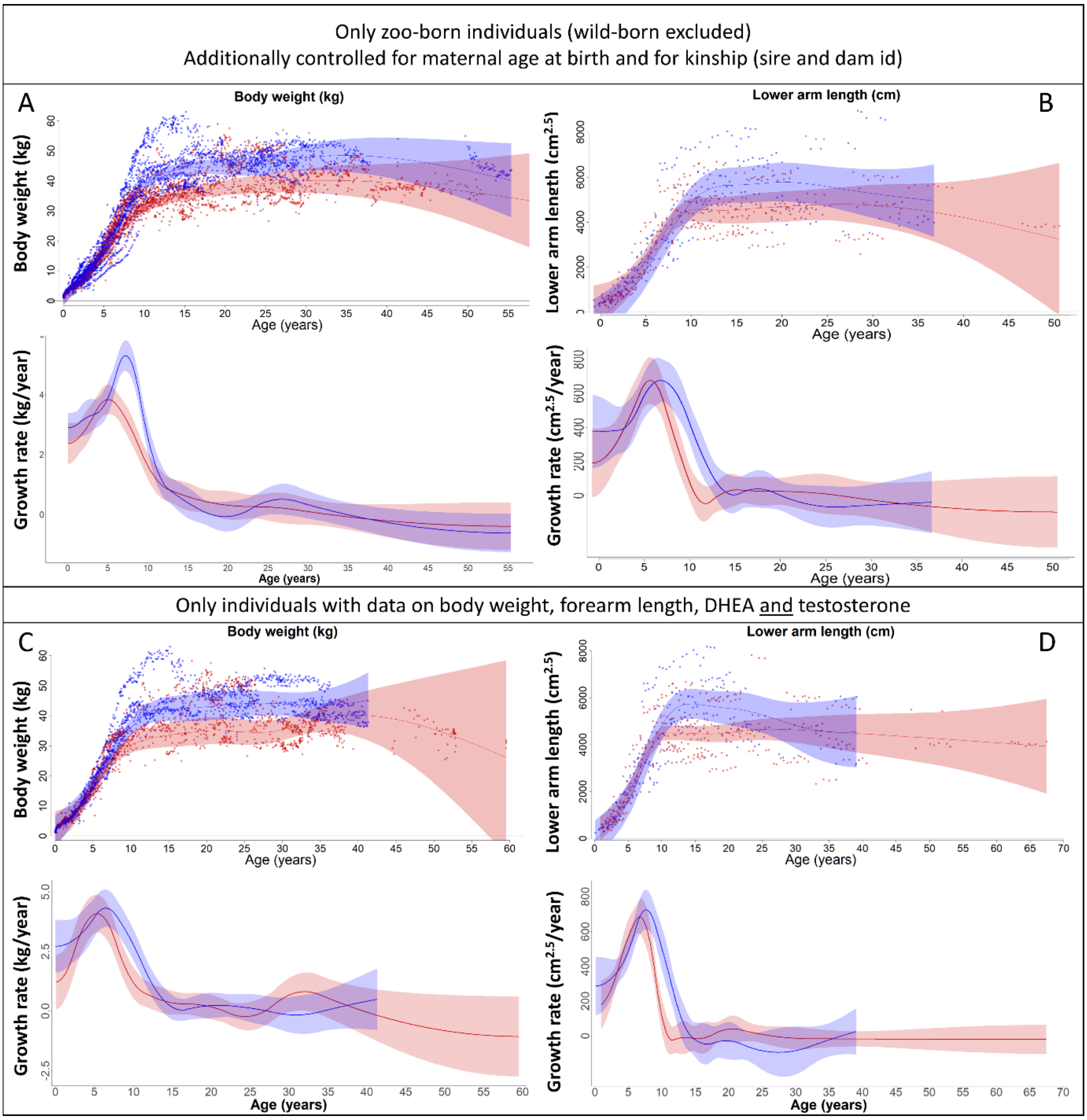


**Figure 2–figure supplement 1**. Our results on body weight (left) and forearm length (right) remained the same if (**A, B**) only zoo-born individuals were considered (which also allowed to additionally control for kinship (dam and sire) and maternal age at birth) or if (**C, D**) only those individuals were considered for which data on body weight, forearm length, DHEA and testosterone were available. Blue = males; red = females. 95% confidence intervals are plotted.

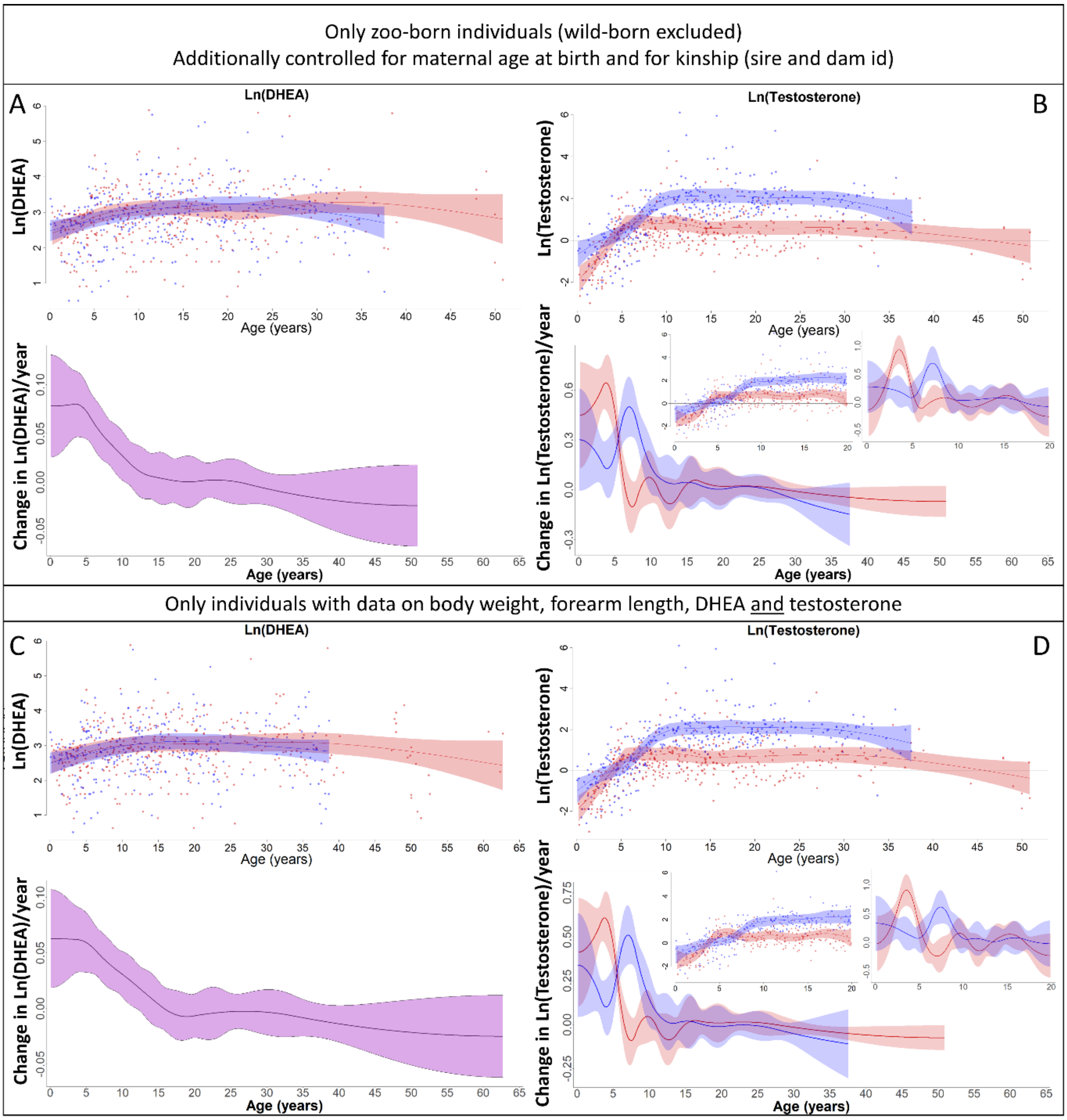


**Figure 3–figure supplement 1.** Our results on DHEA (left) and Testosterone (right) remained the same if (**A,B**) only zoo-born individuals were considered (which also allowed to additionally control for kinship (dam and sire) and maternal age at birth) or if (**C,D**) only those individuals were considered for which data on body weight, forearm length, DHEA and testosterone were available. Blue = males, red = females. 95% confidence intervals are plotted.

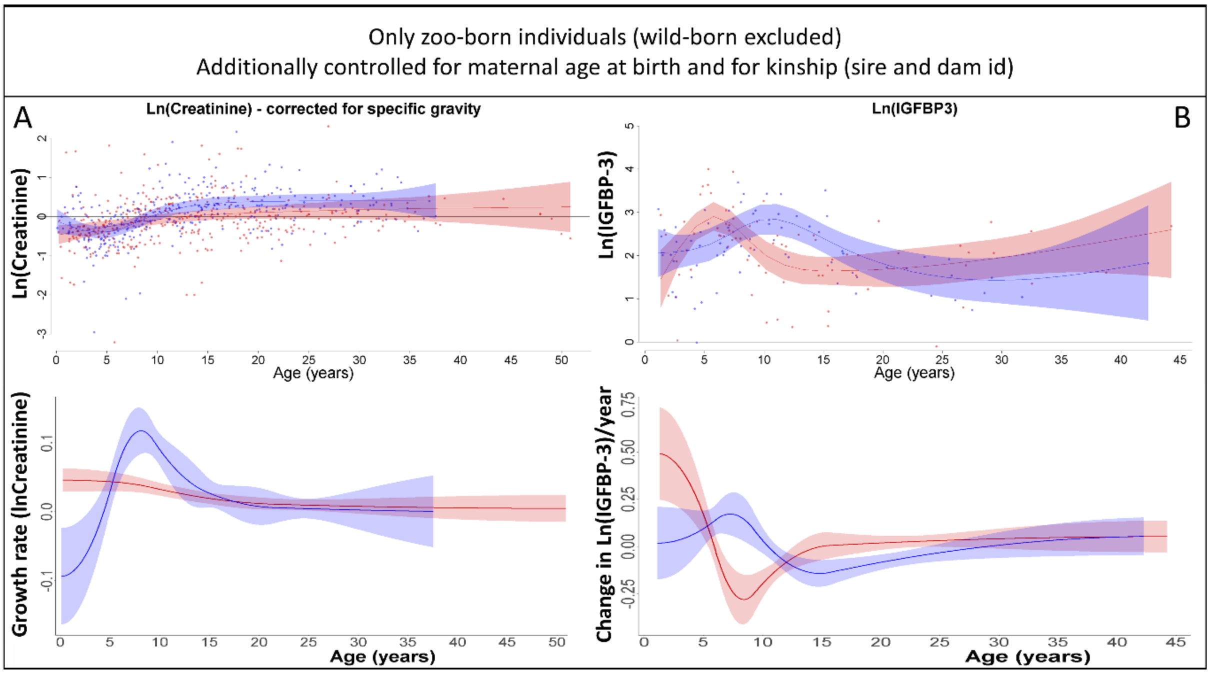


**Figure 3–figure supplement 2.** Our results on creatinine (**A**) and IGFBP-3 (**B**) remained the same if only zoo-born individuals were considered (which also allowed to additionally control for kinship (dam and sire) and maternal age at birth). Blue = males, red = females. 95% confidence intervals are plotted.

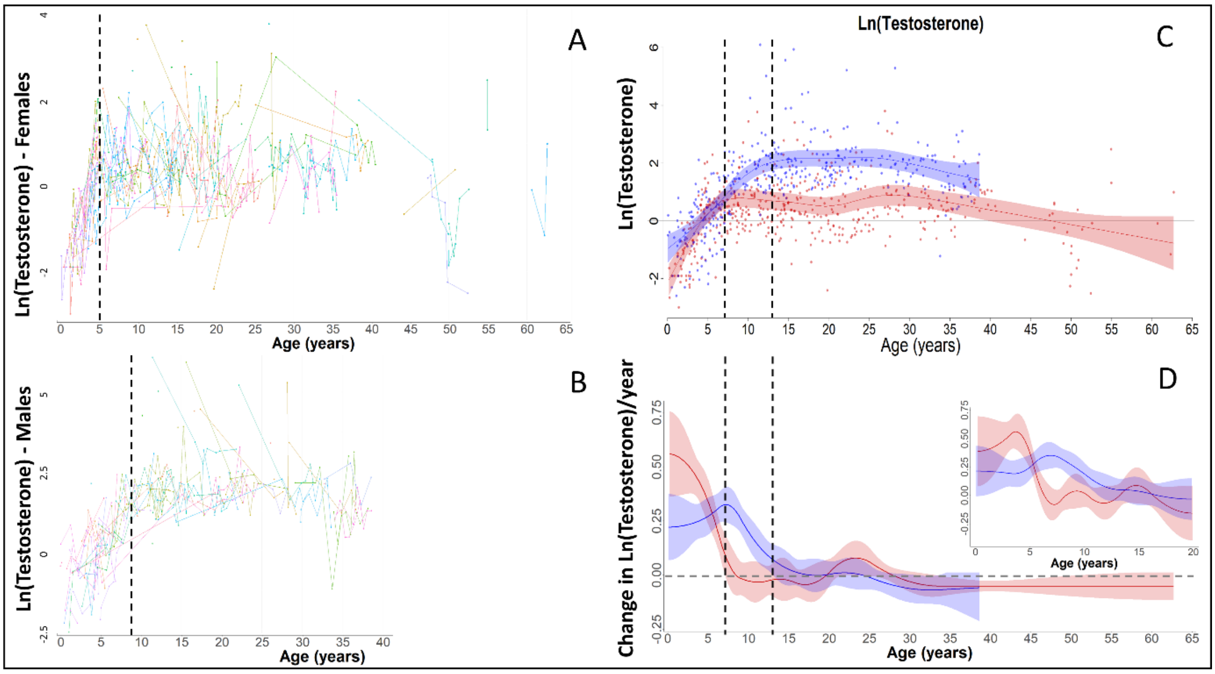


**Figure 3–figure supplement 3.** In our raw data, females and males showed a fast increase in testosterone, as also previously shown for bonobos (37), with females (red) reaching maximal levels around the age of five, and males (blue) around the age of nine years. However, this fast change was not appropriately modelled by our GAMM models if the "automatic" estimation of the smoothing parameters was used, leading to strong oversmoothing and much later ages at which maximal levels were attained, and at which the increase in testosterone levels ceased, in both sexes. Therefore, we applied a fixed smoothing parameter to our testosterone GAMMs of sp = 1, which allowed for higher "wiggliness" and thus faster changes, and which solved the issue.

**Table 2 – table supplement 1. Evidence of (adolescent) length growth spurts (GS) from published literature using linear length growth.** Same table as Table 2 in main text, but additionally with literature on markers of adolescence. Changes in length growth rate (left) - Measures of linear length growth are taken of: Body length or height = B, Crown-rump/Shoulder-rump/Anterior trunk length = CR/SR/AT, Lower/Upper/Full arm length = LA/UA/A, Thigh/Tibia/Leg length = TH/TI/L. Methods: in zoos = direct measurements, in wild populations = photogrammetry, except on *Macaca ochreata* (direct on trapped animals). Growth rate acceleration can be seen as proof of a GS, but taking into account scale-correction, a GS is also very likely in case of a period with constant linear length growth rate, and would be possible in cases of just a slowdown in deceleration. Markers of adolescence (right, often different study population): Testes size growth = TS, Rise in testosterone levels = TL, Menarche = M, First swelling/cycle/ovulation = S/C/O; Timing compared to GS: aligned = a, preceding = p, later (following) = l; m = male, f = female, w = wild, z = zoo .

| Species (w/z) | Changes in length growth rate | | | | | | | Markers of adolescence | | |
|---|---|---|---|---|---|---|---|---|---|---|
| | Acceleration | Constant (plateau) | Slowdown in deceleration | No slowdown in deceleration | Aligned with weight-GS | Comments | Publication | Males | Females | Publication |
| *Macaca assamensis* (w) | | | m + f (LA) | | Not available | Acceleration if scale-corrected | (Anzà et al., 2022; Berghänel et al., 2015) | / | / | / |
| *Macaca fuscata* (z) | m + f (B), m (AT, UA) | m (TH, L) | m (LA), f (LA | f (UA) | Yes (little earlier) | | (Hamada, 1994; Hamada et al., 1999; Hamada and Yamamoto, 2010) | TS (l) | M (a) | (Hamada et al., 1999) |
| *Macaca nemestrina* (z) | m + f (AT, CR, A, LA, L) | | | | Yes | | (Nishikawa, 1985; Tarrant, 1975) | / | S (p) | |
| *Macaca arctoides* (z) | m (CR) | | | | Yes | Few individuals | (Faucheux et al., 1978) | TS (a) | / | (Nieuwenhuijsen et al., 1987) |
| *Macaca mulatta* (z) | (B, TI)[1] | m + f (CR) | | | Yes (little earlier) | [1]Unknown sex, few individuals | (Tanner et al., 1990; van Wagenen and Catchpole, 1956) | / | S (a), M (l) | (Tanner et al., 1990) |
| *Macaca ochreata* (w) | m + f (CR) | | | | Yes | | (Schillaci and Stallmann, 2005) | / | / | / |
| *Theropithecus gelada* (w) | m + f (SR) | | | | Not available | | (Lu et al., 2016) | TL (l) | pigmentation of bare area (a) | (Beehner et al., 2009; Matthews, 1956) |
| *Papio anubis* (z) | m (CRL) | m + f (A) | m (TH), f (CRL, TH) | | Yes (little earlier) | | (Leigh, 2009) | TS, TL, IGF1, IGFBP3 (all a) | C (a) | (Bernstein et al., 2013, 2008; Mueller, 2005; Owens, 1976) |
| *Papio hamadryas* (z) | m (CR) | | | f (CR) | Yes (m) | Coarse data | (Crawford et al., 1997) | TS, TL, IGF1, IGFBP3 (all a) | C (a) | (Bernstein et al., 2013; Mueller, 2005) |
| *Mandrillus sphinx* (z) | | m + f (CR) | | | Yes | | (Setchell et al., 2001) | TS (a), TL (l) | S (a) | (Setchell and Dixson, 2002; Wickings and Dixson, 1992) |
| *Pan troglodytes* (z) | | | m + f (B) | | Yes | | (Hamada and Udono, 2002) | TS & TL (a) | M (l) | (Anestis, 2006; Coe et al., 1979; Kraemer et al., 1982) |
| *Pongo pygmaeus* (z) | m + f (B, LA) | | | | Yes | 2 individuals | (Vančatová et al., 1999) | Highly variable | M (occurs at 5-12yrs) | (Maggioncalda and Sapolsky, 2002; Markham, 1990) |
| *Gorilla beringei beringei* (w) | | | m (B) | m (UA), f (B, UA) | Not available | | (Galbany et al., 2017) | / | / | / |

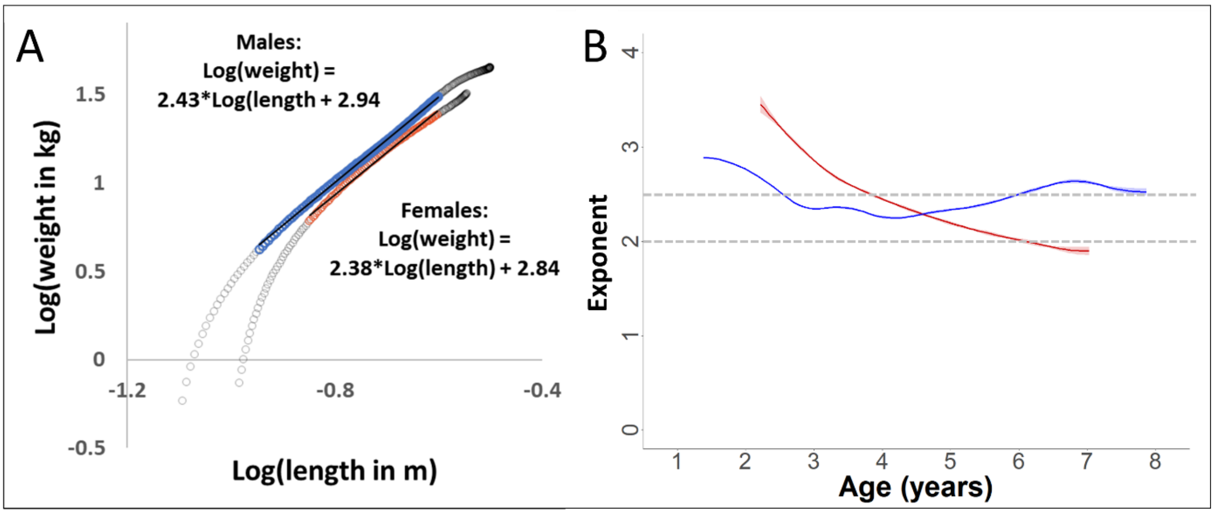


**Appendix 1.** Validation of the scaling relationship between forearm length and body weight during growth in our study. Since weight and length were not measured pairwise, we used the fitted values from the respective GAMM models. The power-exponent of the relationship is calculated as the linear slope of the log-log plot. (**A**) Log-log plot of weight over length, with data from the growth period (range 0-10 years of age in females and 0-14 years in males, grey circles). During the period of a linear relationship (colored, blue=males and red=females), the slope (and thus the exponent of the relationship between the non-transformed variables) was 2.4 for both sexes. (**B**) For the same data used for slope calculation in (**A**), but with dynamic slope calculation using the first derivative of the GAM-smooth. The exponent for males was relatively stable around 2.5 during this age period, whereas the value decreased in females, reaching about 2.0 towards the end of the growth spurt (see results section, **Fig. 2-4**).
